# A novel *Leishmania infantum* reference strain for gene editing and the study of visceral leishmaniasis

**DOI:** 10.1101/2025.02.18.638795

**Authors:** Rokhaya Thiam, Maria Santana Ceballos, Tom Beneke, Nada Kuk, Grégoire Pasquier, Lucien Crobu, Daniel C Jeffares, Baptiste Vergnes, Bridlin Barckmann, Yvon Sterkers

**Affiliations:** University of Montpellier, CNRS, IRD, Academic Hospital (CHU) of Montpellier, MiVEGEC, Montpellier, France; Department of Cell and Developmental Biology, Biocentre, University of Würzburg, Germany; Department of Biology and York Biomedical Research Institute, University of York, United Kingdom

## Abstract

Parasites of the *Leishmania donovani* complex are responsible for visceral leishmaniasis, a vector-borne disease transmitted through the bite of female phlebotomine sand flies. As well as the human hosts, these parasites infect many mammals which can serve as reservoirs. Dogs are particularly important reservoirs in Europe. Transmission is widespread across Asia, Africa, the Americas, and the Mediterranean basin, including South of France. Visceral leishmaniasis poses a fatal threat if left untreated. Research into the pathophysiology of this neglected disease is of prime importance, as is the development of new drugs. In this study, we evaluated the growth, differentiation, and macrophage infectivity of four *L. donovani* complex strains and identified *L. infantum* S9F1 (MHOM/MA/67/ITMAP263, clone S9F1) as a well-adapted strain for genetic engineering studies. We present here the genome sequence and annotation of *L infantum* S9F1 T7 Cas9, providing the scientific community with easy access to its genomic information. The data has been integrated into the LeishGEdit online resource to support primer design for CRISPR-Cas9 experiments. We now aim to make this strain widely available to foster pathogenesis studies of visceral leishmaniasis.

**AUTHOR SUMMARY:** Visceral leishmaniasis is a disease caused by parasites of the *Leishmania donovani* complex. These parasites are spread to humans and animals through the bites of sand flies, and this disease affects millions of people worldwide, particularly in regions such as the Americas, Asia, Africa, and the Mediterranean basin. If left untreated, it can be fatal. Researchers need to study the biology of the parasite that causes the disease to better understand how it develops and progresses. In this study, we identified a *L. infantum* strain that is amenable to genetic modification in the laboratory and may serve as a representative model of the causative agent of visceral leishmaniasis. We tested two CRISPR-Cas9 strategies on this strain, re-sequenced and annotated its genome, and made the data available on the LeishGEdit website. By sharing this strain with the research community, we aim to support further studies on the pathogenesis of visceral leishmaniasis.

## INTRODUCTION

Leishmaniasis is a vector-borne disease caused by approximately twenty species of flagellated protozoa belonging to the genus *Leishmania* (1). It is transmitted to humans through the bite of infected female sandflies (1). The parasite undergoes two distinct evolutionary stages during its life cycle (2). The promastigote stage is an extracellular, motile form facilitated by its flagellum and is typically found in the sand fly’s intestine. In contrast, the amastigote stage is intracellular, residing primarily within the parasitophorous vacuole of macrophages in the mammalian host (2). *Leishmania* spp., especially those causing the severe forms, impose a global health and economic burden. Leishmaniasis is endemic in over 98 countries, with approximately 350 million people at risk (3). There are an estimated 12 million cases of infection, with 700,000 to 1 million new cases reported annually, resulting in 20,000 to 30,000 deaths per year (3,4). Visceral leishmaniasis, also known as kala-azar in India, is fatal if left untreated. It is characterized by irregular fever, weight loss, hepatosplenomegaly, and anemia (1,5). The main causative agents of visceral leishmaniasis are species within the *Leishmania donovani* complex, including *L. donovani* in Asia and Eastern Africa, and *L. infantum* in Latin America, Central Asia, Eastern and Northern Africa, and the Mediterranean region (1,4,6–8).

As in many fields, the introduction of CRISPR-Cas9 has been a major revolution in the study of leishmaniasis. Since its implementation in 2015 (9), several strategies have been used (10), in particular a PCR-based strategy hereafter LeishGEdit system (11) and a Ribonucleoprotein-based strategy (12,13). To promote studies on the pathogenesis of visceral leishmaniasis, we wanted to make available to the community a strain of *L. donovani* complex for which we have re-sequenced the genome and tested the CRISPR-PCR and RNP-based strategies.

In order to identify an appropriate strain for *in vitro* studies, we screened four strains of the *L. donovani* complex. We examined, cell growth, capacity to differentiate into amastigotes, and ability to infect macrophages to identify *L. infantum* S9F1 (MHOM/MA/67/ITMAP263, clone S9F1) as well adapted for future work. We adapted the LeishGEdit system to this strain by transfecting an episomal which encodes Cas9 and T7 polymerase, while specific sgRNA and donor DNAs are produced by PCRs (11). As proof of concept, we successfully tagged and deleted the flagellar protein PF16. We also implemented and tested a ribonucleoprotein complex strategy for CRISPR-Cas9 editing, where Cas9 protein and guide RNA are produced *in vitro* and complexed prior to transfection (12,13). In total, we succeeded in identifying a well-adapted strain for genetic engineering studies, exhibiting high growth rates in axenic amastigotes and promastigotes, as well as a satisfactory parasitic index in macrophage infections.

## MATERIAL AND METHODS

### Strains

*L. infantum* S9F1 (MHOM/MA/67/ITMAP263 clone S9F1, obtained from a human patient in Morocco in 1967 and passed in a mouse, hereafter S9F1), *L. infantum* LEM6696 (MCAN/ES/98/LLM-877, JPCM5, obtained from a dog in Spain in 1998), *L. donovani* LEM703 (MHOM/IN/80/DD8; ATCC Cat. No. 50212, obtained from a human patient in India in 1980), *L. donovani* Ld1S-Bob (a cloned line derived from the isolate MHOM/SD/62/1S-CL2D, obtained from a human patient in Sudan in 1962), *L. mexicana* (MNYC/BZ/62/M379) originally isolated from a Sumichrast’s vesper rat (*Nyctomys sumichrasti*) in 1962 in Belize and *L. mexicana* T7 Cas9 derived from the same genetic background by transfection of the pTB007 plasmid (11).

### Promastigote culture and growth curve determination

Promastigotes were grown by incubation of parasites cultures at 27°C in HOMEM medium (14) supplemented with 10% fetal bovine serum (Gibco, A3160801) and hemin at a final concentration of 6 µg/mL. For growth curves, cultures were monitored daily by counting parasites with a Cell Drop automated cell counter (DeNovix). When the parasites were in the exponential phase, the cultures were diluted in fresh medium.

### Transition to axenic amastigote

Promastigotes were grown to the stationary phase then centrifuged at 1000g for 5 min. The pellet was resuspended in Schneider medium pH 5.5 supplemented with 20% fetal bovine serum. Parasites were incubated at 32°C with 5% of CO_2_ (15). Promastigote to amastigote transition takes three to five days, all experiments on amastigotes were performed after day 7.

### *In vitro* macrophage infections

THP-1 cells were cultivated in RPMI 1640+GlutaMAX, 25mM HEPES, supplemented with MEM non-essential amino acids (Thermofisher # 72400047), 100 mM sodium pyruvate, penicillin (10,000U/mL), streptomycin (10,000 mg/mL) and 10% fetal bovine serum. Each well of a sterile 16-well Lab-Tek chamber slide was filled with 200µL of THP-1 cells (10⁵ cells/mL) in medium containing a final concentration of 100 ng/mL phorbol myristate acetate (PMA). Cells were incubated at 37°C with 5% CO_2_ for 48h to adhere to the coverslips. A wash with non-supplemented RPMI medium was performed to remove non-adherent cells. Parasites were washed three times with non-supplemented RPMI. Infections of the cultures were performed with either axenic amastigotes or promastigotes; in both cases the multiplicity of infection (MOI) was 5:1. After 4h of infection, the cultures were washed with non-supplemented RPMI to remove extracellular parasites and maintained in culture for 24h at 32°C with 5% CO_2_ to allow parasites to replicate and then 24h at 37°C with 5% CO_2_. The cells were then fixed with methanol and stained with Giemsa stain. Two hundred macrophages were counted per condition. Two parameters were considered: the percentage of infected THP-1 and the parasitic index 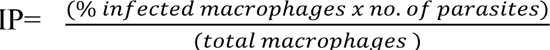 as described elsewhere (16).

### Viability test and drug delectable marker sensitivity testing

To check the sensitivity of the strain and assess the half maximal inhibitory concentration (IC50) of the main selectable markers used for genetic engineering, methylthiazolyldiphenyl-tetrazolium bromide (MTT) tests MTT assays were performed; the yellow MTT dye is reduced by mitochondrial enzymes in living cells to form blue formazan, detected by spectrophotometry. We tested hygromycin, blasticidin, geneticin, puromycin, nourseothricine (NTC) and phleomycin (all from InvivoGen). Briefly, S9F1 promastigotes in the exponential phase of growing were diluted to a parasite concentration of 2 x 10^6^ cells/mL. In 96 well plates, serial dilutions of the drugs were performed, the parasites were added, and the plate incubated at 27°C for 72 hours. MTT (Sigma-Aldrich CAS 298-93-1) was added to the culture according to the manufacturer specifications. The plates were then incubated in the dark at 37°C for 4h. Optical density was read between 550 and 690 nm with a spectrophotometer. The percentage of viability was determined by calculating the ratio between the OD obtained for each concentration tested and the OD obtained with untreated parasites.

### Cell motility assay

Five µL from a cell culture at a density of 5×10^6^ cells/mL were deposited on a glass slide as described elsewhere (17). Thirty-second videos were immediately recorded at five frames per measured using the open-source Fiji plugin TrackMate (18). At least five videos for each strain were recorded and analyzed.

### Episomal expression of Cas9

Cells were transfected using the Amaxa Nucleofector^TM^ technology. As described elsewhere (19), 3×10^6^ parasites in exponential phase were transfected with 20 µg of the plasmid (pTB007) This plasmid contains the S*treptococcus pyogenes* Cas9 nuclease gene (*SpCas9*), T7 RNA polymerase gene and a hygromycin resistance gene (11). The transfection was performed using a transfection buffer containing 200 mM Na₂HPO₄, 70 mM NaH₂PO₄, 15 mM KCl, 150 mM CaCl₂, and 150 mM HEPES (pH 7.3) and slot cuvettes. Cells were transfected using the X-001 program of the Amaxa Nucleofector IIb (Lonza Cologne AG, Germany). The transfected cells were immediately transferred to 25 cm^2^ flasks containing 5 mL of pre-warmed medium, and then incubated at 26°C. On the next day, 50 µg/mL hygromycin was added to the culture medium and stable Cas9 transfectants were obtained between 10 and 14 days after transfection.

### Nuclear staining and indirect immunofluorescence assay

Cells were fixed in 4% paraformaldehyde and air-dried on microscope immunofluorescence slides. Slides were treated with 0.2% Triton ×100 in PBS to permeabilize membranes and saturated with 1% BSA in PBS. Cells were incubated with either anti-mNeonGreen (1:1000 dilution, mNG, 32F6-Chromatek), anti-mCherry (1:1000 dilution, GTX128508 Gene Tex) and anti-HA (1:200 dilution, Sigma). After three washes with 1% BSA in PBS, the slides were incubated with anti-IgG specific secondary antibodies, conjugated with Alexa Fluor 546 (Molecular Probes®, ref. A-11010) or Alexa Fluor 488 (Molecular Probes®, ref. A-11001) diluted to 1:1500. The slides were washed with PBS, and the DNA was stained with Hoechst 33342 (Thermo Scientific) (1:5000 dilution) and mounted with Vectashield (Vector Laboratories®).

### Western blot analysis

Protein extracts from 4×10^6^ cells, diluted in Laemmli 2X, were loaded per well, separated by SDS-PAGE, and transferred onto PVDF membranes (Hybond-P, Amersham). Primary antibody against Cas9 (Diagenode) was used at a 1:2500 dilution and anti-LACK (20) was used at a 1:3000 dilution as a loading control and for normalization. The membranes were incubated for one hour at room temperature with alkaline phosphatase conjugated secondary mouse or rabbit antibodies (Promega) at 1:3000 dilution respectively and revealed with 5-bromo-4-chloro-3-indolyl phosphate/nitro blue tetrazolium (BCIP/NBT; Promega) according to manufacturer’s instructions.

### CRISPR-Cas9 edition

#### PCR-based method for tagging and knock-out

The deletion, 3’ mNG or 3’ mCherry gene tagging of the PF16 gene (PF16: LINF_200019300, LinJ.20.1450) (21) were performed using CRISPR-Cas9 as previously described (11). The online primer design tool LeishGEdit (www.LeishGEdit.net, last access January, 28^th^ 2025) was used to design primers, using the JPCM5 genome (7). For the gene deletion of PF16, primers were Upstream forward primer: CCCTCGACACAGACACAAACTACACGGGCAgtataatgcagacctgctgc and Downstream reverse primer: CGAGCAGCGTGAGTTGGCGTGGCCGTGCCGccaatttgagagacctgtgc for the donor DNA. In lower case, the specific primers of the pPLOT plasmids and in upper case the specific homology regions of the target gene. For tagging, the tag is amplified using a specific pPLOT plasmid.They were gaaattaatacgactcactataggCACGTATCAGCGCTACACAGgttttagagctagaaatagc and gaaattaatacgactcactataggGCTGATGCTCAGCCGCTATTgttttagagctagaaatagc for the 5’ and 3’ sgRNA primer. For 3’ tagging, primers were: gaaattaatacgactcactataggGCTGATGCTCAGCCGCTATTgttttagagctagaaatagc for 3’ sgRNA primer, and for donor DNA, the downstream forward primer was AAGATCGAGAACTACCACGTGCAGCAGCACggttctggtagtggttccgg, and the downstream reverse primer: CGAGCAGCGTGAGTTGGCGTGGCCGTGCCGccaatttgagagacctgtgc. In lower case are the sequences of the T7 promoter on one side and the constant structure of the sgRNA scaffold on the other, in upper case the seed oriented 5‘ to 3’ (note that the 3’ PAM is not in the construct).

#### Ribonucleoprotein-based (RNP)-based CRISPR-Cas9 method

For the 3’ gene tagging using the RNP-based strategy, the seed was also designed on LeishGedit website targeting PF16: LINF_200019300, LinJ.20.1450 in the JPCM5 reference genome. As compared to the PCR-based strategy, only the nucleotides indicated in capital letters in the sgRNA primer sequence were used (GCTGATGCTCAGCCGCTATT). The sgRNA was ordered from Integrated DNA Technologies (IDT) in Leuven, Belgium, which synthesized the crRNA and tracRNA (trans-activating CRISPR RNA) separately. Here, since the 3’ gene tagging was marker free, only the pPLOT downstream forward primer designed on LeishGedit for the donor DNA could be used. New downstream reverse primers were designed: Forward (AAGATCGAGAACTACCACGTGCAGCAGCACggttctggtagtggttccgg) and Reverse (CGAGCAGCGTGAGTTGGCGTGGCCGTGCCG<u>tta</u>actacccgatcctgatccagat). These primers include a stop codon in phase with the reading frame, highlighted here (TTA reverse complementary form of TAA).

#### Transfection of parasites

Cells were transfected using the Amaxa Nucleofector™ technology, program X001. Approximately 5×10^6^ parasites in logarithmic phase were transfected with 20 µg of donor DNA and sgRNA. For RNP editing the sgRNA was generated by an equimolar mix of 200 µM of crRNA and tracRNA that was heated at 94°C for 5 minutes then kept at −20°C for further usage. For editing 50 µg of Cas9 protein from IDT were complexed at room temperature with the prepared sgRNA according to the manufacturer protocol and then transfected. In knockout experiments, antibiotics were added in the culture medium at day 1.

#### Whole genome sequencing

DNA extraction was performed on S9F1 T7 Cas9 cells, using the Qiagen DNeasy Blood and Tissue Kit according to the manufacturer’s instructions. The sequencing library was prepared using the Nextera XT DNA Library Preparation Kit (96 samples), following the protocol in the Nextera XT DNA Library Prep Kit Reference Guide (15031942 v03). The whole genome sequencing was performed generating 150 nt paired-end reads on a NovaSeq X (Illumina). The resulting raw sequence data were analyzed, with FastQC v0.11.9 (https://www.bioinformatics.babraham.ac.uk/projects/fastqc/, last access January, 28^th^ 2025) to evaluate sequence quality. Cutadapt v4.0 (23) as used to remove adapter sequences and Trimmomatic v0.39 (24) employed to trim low-quality bases. The reads were then deduplicated using clumpify from BBTools v39. This process produced 17,800,790 filtered, non-redundant reads, which were subsequently used for downstream analyses.

To generate the genome sequence for the S9F1 T7 Cas9 strain, we used the JPCM5 reference genome from TriTrypDB (https://tritrypdb.org/tritrypdb/app, last accessed January 28, 2025) as a S9F1 T7 Cas9 whole-genome sequencing (WGS) reads. To do so filtered, non-redundant reads were first aligned to the JPCM5 genome using BWA-MEM2 v2.2.1 (25) resulting in an average genome coverage of 76. The resulting BAM files were converted, sorted, and indexed with Samtools (26). Next, we performed the initial round of Pilon v1.23 polishing to generate the predicted S9F1 T7 Cas9 genome sequence. To further improve accuracy, we applied an additional round of Pilon polishing to the new genome sequence.

To generate a gene annotation for the new genome sequence we used the tool Liftoff v1.6.3 which uses the genome annotation of a closely related genome in gff file format, in our case JPCM5, and maps it to the new genome sequence by aligning each gene from the reference genome to the target genome and finding the mapping that provides highest sequence identity while preserving the gene structure. LiftoffTools (27) was then used to compare the two genomes and their annotations for gene sequence variants, synteny, and gene copy number changes.

#### Incorporation of the S9F1 T7 Cas9 *L. infantum* genome on LeishGEdit.net

LeishGEdit primers for the polished S9F1 T7 Cas9 genome were designed using a standalone bash script (28) and are available under www.leishgedit.net. Primers for geneIDs can be queried using “LINS9F1C” instead of LINF (e.g. LINS9F1C_010005000 instead of LINF_010005000). A 17-nt DNA barcoded upstream forward primer can be also obtained (28), enabling the generation of barcoded mutant pools for future studies.

#### Statistical analysis

For the motility assays, graphs and statistical tests were performed using R. The software was from R Core Team (2024). *R: A language and environment for statistical computing.* R Foundation for Statistical Computing, Vienna, Austria. Available at: https://www.R-project.org/. The package used was from Wickham H (2016): *ggplot2: Elegant Graphics for Data Analysis.* Springer-Verlag New York. Available at: https://ggplot2.tidyverse.org. For all other figures, graphical illustrations and statistical analyses were conducted using Prism software (GraphPad Software, Version 8).

## RESULTS AND DISCUSSION

### S9F1 identified as a promising strain for *in vitro* studies

We evaluated four strains of the *L. donovani* complex: *L. infantum* JPCM5 (MCAN/ES/98/LLM-877, LEM6696), *L. infantum* S9F1 (MHOM/MA/67/ITMAP263 clone S9F1), *L. donovani* Ld1S-Bob, and *L. donovani* LEM703 (MHOM/IN/80/DD8; ATCC Cat. No. 50212) which are routinely cultured in our laboratory. Additionally, we included as a control strain *L. mexicana* T7 Cas9 from the genetic background of MHOM/GT/2001/U1103. The latter being the reference strain utilized for functional genome-wide screens (11,29). Initially, we assessed the cell growth of the promastigote forms of these strains and observed similar growth rates for all strains (Fig. 1A). Subsequently, we examined their capacity to differentiate and to grow into axenic amastigotes and their ability to infect macrophages. All strains differentiated into axenic amastigotes. *L. donovani* Ld1S-Bob and S9F1 grew to reach a maximum cell concentration of 2 x 10^7^cells/mL, while *L. mexicana* T7 Cas9 grew to a maximum of 2 x 10^8^ cells/mL. *L. donovani* LEM703 exhibited lower growth rates compared to the other strains and in our hands, *L. infantum* LEM6696 (JPCM5) did not even grow at all as axenic amastigotes (Fig. 1B). Regarding *in vitro* infectivity, S9F1 had the highest percentage of infecting THP-1 derived macrophages; particularly when amastigotes were used to seed the infection and the highest parasitic index as compared to other strains (Fig. 1C-D). We were surprised by the results obtained with LEM6696, which has the same genetic background as JPCM5, because this strain has been shown elsewhere to efficiently transform into amastigotes and proliferate inside monocytes (30). The loss of proliferative capacity and macrophage infectivity of LEM6696 could be attributed to extensive passaging, resulting in the loss of virulence. Similar assumptions could be made regarding LEM703, which also exhibited reduced macrophage infectivity *in vitro*. One of the limitations of our study is that not all strains had the same number of passages since the last inoculation to the animal. Consequently, these results should not be generalized, and it should not be concluded that one strain is intrinsically more virulent than another. It is important to note that virulence may need to be restored periodically by inoculation in an animal model and that not all strains necessarily lose their virulence at the same rate, which was not tested here.

**Figure 1:**
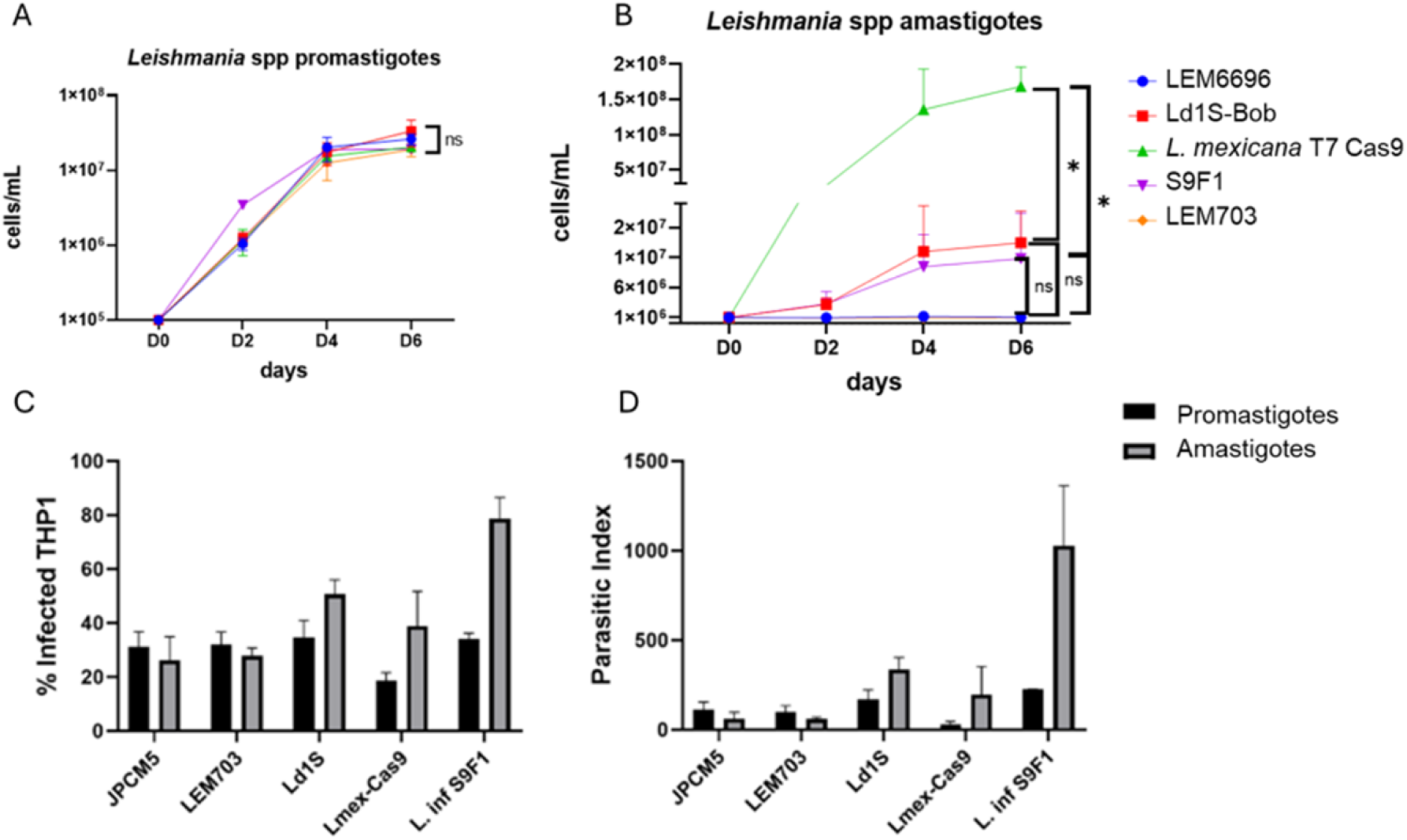
Characterization of *in vitro* growth and macrophage infectivity (A) *L. infantum* JPCM5 (MCAN/ES/98/LLM-877, LEM6696), *L. infantum* S9F1 (MHOM/MA/67/ITMAP263 clone S9F1), *L. donovani* Ld1S-Bob, and *L. donovani* LEM703 (MHOM/IN/80/DD8) and *L. mexicana* T7 Cas9 promastigote forms in HOMEM incubated at 27°C. (B) Growth curves of amastigote forms cultivated in Schneider medium pH 5.5 incubated at 32°C. Equal number of *L. donovani* complex strains and *L. mexicana* T7 Cas9 promastigotes (10^5^cells/mL) or the amastigote forms (10^6^cells/mL) were inoculated in flask containing 5 mL of HOMEM or Schneider medium. The parasite growth was monitored by using a cell counter once a day for a total of 6 days. Statistical tests used: One-way ANOVA followed by Tukey’s test, ns: not significant, *= p-value < 0.05. (C-D) Infection of THP-1 derived macrophages with promastigote (black histograms) and amastigote forms (grey histograms) (MOI 5:1). The percentage of infected macrophages (C) and the parasitic index (D) were determined by microscopy, counting 200 infected and non-infected THP-1 cells and the number of amastigotes within infected macrophages. Three independent experiments were performed.

In summary, we consider that among the strains tested, S9F1 exhibited the highest parasite index and demonstrated good growth rates in both the promastigote and amastigote stages, making it a most suitable strain for *in vitro* studies of a causative agent of visceral leishmaniasis, based on these experiments. We therefore decided to further characterize this strain.

### Characterization of *L. infantum* S9F1 cell cycle

Here we aimed to characterize the cell cycle progression of S9F1 cells and to compare it with that of *L. mexicana* T7 Cas9. *Leishmania* species have an unconventional form of mitosis known as closed mitosis, in which the nuclear envelope does not disassemble during cell division (31,32). *Leishmania* spp, like other trypanosomatids are characterized by a single mitochondrion containing the mitochondrial DNA, known as “kinetoplast DNA”. The latter is a highly complex concatenated network of mini- and maxi-circles visible under the microscope after DNA staining (33). Nuclear and mitochondrial cycles are independent yet coordinated (34). Cell cycle stages can therefore be determined by analyzing the number of kinetoplasts, nuclei, and flagella present in a cell, hereafter called Nuclear Kinetoplast Flagellum (NK) pattern. During the interphase, parasites one kinetoplast, one nucleus, and one flagellum (1K1N1F). Cell division involves first the duplication and elongation of the flagellum (1K1N2F). During cytokinesis, parasites present the pattern 2K2N2F (35,36). Here NK pattern analysis showed the same chronology of division of the different organelles (Kinetoplast (K), Nucleus (N), Flagellum (F) within the cell in *L. mexicana* T7 Cas9. and S9F1 (Fig. 2A). More specifically, the analysis of the NK pattern showed that there were no statistically significant changes in the different cell cycle stages between S9F1 and *L. mexicana* T7 Cas9 (Fig. 2B). Measurements revealed some morphological differences between S9F1 and *L. mexicana* T7 Cas9. S9F1 presents a cell body and a flagellum length shorter than *L. mexicana* T7 Cas9, regardless of whether the parasite is in the exponential or stationary phase (Fig. 2C).

**Figure 2:**
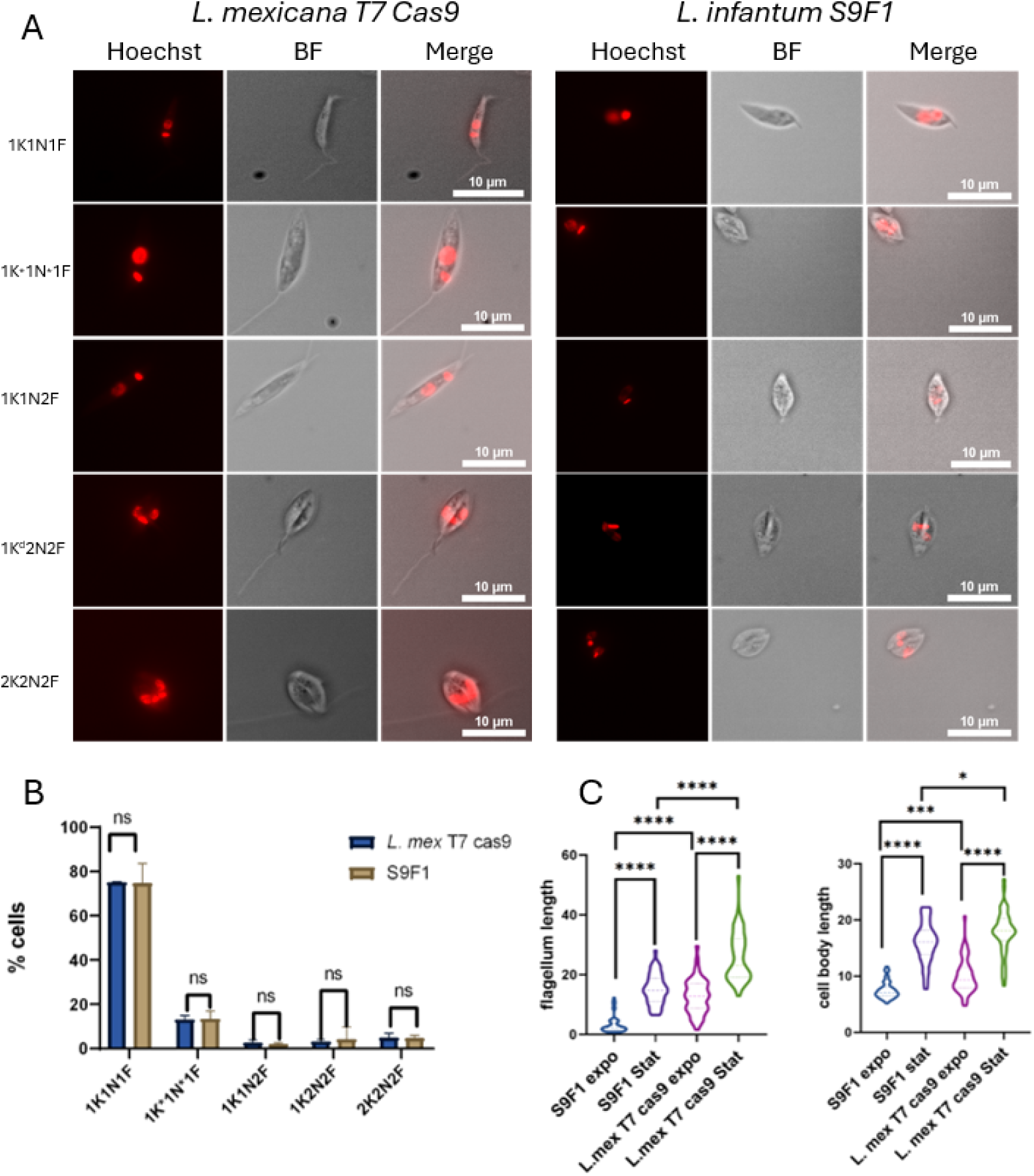
Analysis of the morphology and the cell cycle of *L. infantum* S9F1 and *L. mexicana* T7 Cas9. (A) Different morphological patterns observed in exponentially growing *L. infantum* S9F1 *vs. L. mexicana* T7 Cas9 promastigote forms by microscopy. (K) represents the kinetoplast, (N) the nucleus, and (F) the flagellum. (B) NK pattern analysis of *L. infantum* S9F1 vs. *L. mexicana* T7 Cas9 promastigote forms. The data were obtained from 600 cells counted from three independent experiments. Statistical test used: Multiple t-test (two-sided), ns: not significant. (C) Comparison of the flagellum length of *L. infantum* S9F1 vs. *L. mexicana* T7 Cas9 promastigote forms in exponential and stationary phases. (left); of the cell body length of *L. infantum* S9F1 *vs L. mexicana* T7 Cas9 in exponential and stationary phases (right). Measurements were done using ImageJ. One-way ANOVA followed by Tukey’s test was used to compare flagellum length and cell body length. Significance levels: **** p-value < 0.0001, ***p-value<0.001 and *p-value<0.05.

### Establishment of S9F1 T7 Cas9

We tested the sensitivity of S9F1 to the six antibiotics most used in genetic engineering studies: hygromycin, blasticidin, geneticin, puromycin, nourseothricin (NTC) and phleomycin. The IC50s we determined using MTT assays for the six selectable markers were comparable to those obtained in other strains and suitable for genetic engineering (Fig. 3A). We then transfected S9F1 with pTB007 that carries the *SpCas9* and the T7 RNA polymerase genes (11). The resulting hygromycin-resistant cells (hereafter referred to as S9F1 T7 Cas9) were recovered after 10-11 days. Immunofluorescence imaging revealed a whole-cell localization of Cas9 in both S9F1 T7 Cas9 and *L. mexicana* T7 Cas9, while the signal was absent in the parental cell line (Fig. 3B). Western blot analysis confirmed the expression of Cas9 in S9F1 T7 Cas9 but also revealed several lower molecular weight bands in addition to the expected 150 kDa band (Fig. 3C). S9F1 T7 Cas9 grew well as promastigote forms and based on the results of the NK pattern, Cas9 expression does not affect the cell cycle of the parasite (Fig. 3D).

**Figure 3:**
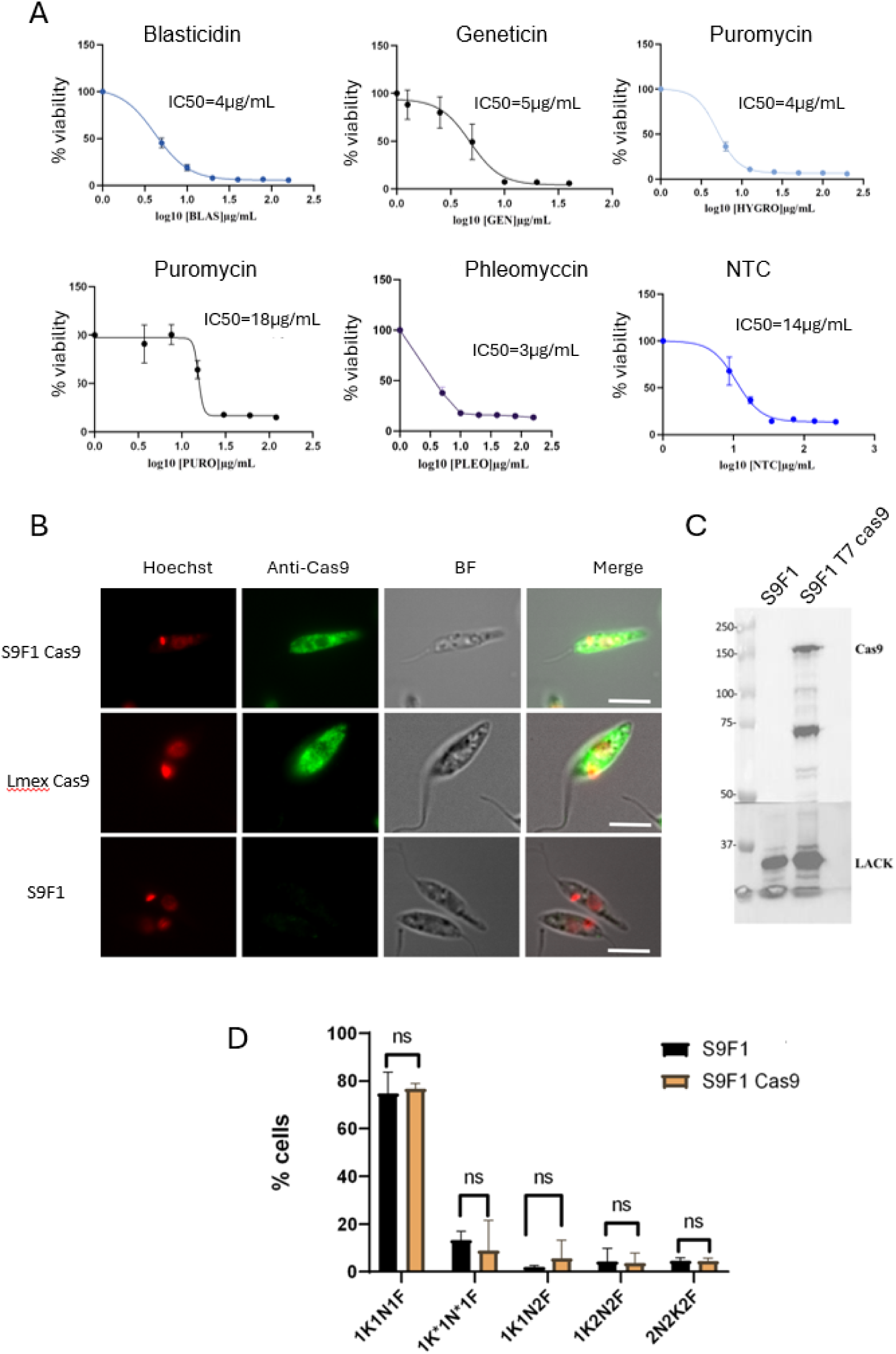
*L. infantum* S9F1 T7 Cas9 characterization (A). IC50s were determined by MTT test for blasticidin, geneticin, hygromycin, puromycin, phleomycin, and Nourseothricin (NTC). The IC50 values, representing the concentration of each drug required to inhibit 50% of parasite viability, were calculated based on the results of an MTT assay. Data points represent the mean ± standard deviation (SD) from three independent biological replicates, each performed in triplicate. (B) Immunofluorescence assay of transfected *L. infantum* S9F1 strain with the vector carrying the Cas9 cassette (pTb007) (top line), *L. mexicana* T7 Cas9 which already expresses Cas9 was used as positive control (middle line) and *L. infantum* S9F1 (bottom line) as negative control. (C) Western Blot analysis of *L. infantum* S9F1 and S9F1 T7 Cas9, expected size of Cas9=150 kDa. LACK is the loading control at 32 kDa. (D) NK pattern analysis of S9F1 and S9F1 T7 Cas9, statistical test used for growth curves and NK pattern: Multiple unpaired t test (two-sided), ns: not significant, ****p value < 0.0001.

### *L. infantum* S9F1 T7 Cas9 genome sequencing

To facilitate genetic engineering on the S9F1T7 Cas9 strain, we re-sequenced its genome. A predicted genome for this strain was generated by ‘polishing’ the reference genome with reads from the S9F1 T7 Cas9 strain using Pilon to introduce single nucleotide polymorphism (SNP) and small insertion/deletion (indel) differences. After this initial round of polishing, an additional round of Pilon polishing was performed to further improve the accuracy of the new genome sequence. The total number of differences observed between the new S9F1 T7 Cas9 genome and the reference genome *L. infantum* JPCM5 (MCAN/ES/98/LLM-877) is 2,469. Out of the 2,469 total changes, 2,237 were identified in the S9F1 T7 Cas9 Pilon assembly when compared to the reference genome. An additional 258 changes were introduced during the second round of Pilon polishing applied to the new assembly. Among these 258 changes, 13 were reversions, where the change initially introduced by pilon in the first round was changed back to the original reference genome sequence. The 2,469 total changes include insertions, deletions, and substitutions (Table 1).

**Table 1:** Summary of genome changes between S9F1-Cas9 and the JPCM5 reference genome both genic and intergenic regions.

| Total number of changes | Number of insertions/deletions/substitutions | Length (nucleotides) | Number |
| --- | --- | --- | --- |
| 2469 changes | 845 insertions | <4 | 823 |
|  |  | ≥4 | 22 |
|  | 758 deletions | <4 | 704 |
|  |  | ≥4 | 54 |
|  | 866 substitutions | >1 | 118 |
|  |  | =1 | 748 |

This table shows the total number of changes, between S9F1 T7 Cas9 and the reference JPCM5, which is the sum of insertions, deletions, and substitutions. The column "Length" specifies the size of these changes, and the column "Number" indicates the count of occurrences for each category.

Thus, the final genome sequence of the S9F1 T7 Cas9 strain contains 2,469 differences from the reference. A subset of reads generated by the sequencing of the S9F1T7 Cas9 strain (∼36229 reads, 0.43% of the aligned reads), mapped to the plasmid used to generate the S9F1 T7 Cas9. No read pairs or reads bridging the plasmid sequence and the genome sequence could be detected and the regions of the plasmid that would be excised after integration into the genome also had a strong read coverage, indicating that the plasmid is present as an episome in the strain. Indeed, the average coverage of the plasmid was 709 which is around 10-fold the read coverage to the reference genome (76 average coverage).

To generate the S9F1 T7 Cas9 genome annotation we used LiftoffTools (27), which uses an annotation from a closely related genome to “lift it over” to another genome. Liftoff aligns genes from the reference genome to the target genome and finds the most likely location. Here, the closely related genome was JPCM5. Liftoff could not map four genes that are in repetitive regions (LINF_100010700, LINF_100010900 encoding GP63 – leishmanolysin, LINF_340023500 and LINF_340034600 corresponding to a putative Amastin-like surface glycoprotein). Given the sequencing method used (no long reads), the mapping method used to generate the predicted S9F1 T7 Cas9 genome, and the repetitive nature of these genes, we cannot conclude that the number of copies of these genes varies between these two strains. These could well be artefacts of the mapping, or the genome assembly via pilon polishing. To study the possible impact of these differences between the reference genome and the S9F1 T7 Cas9 genome we used LiftoffTools (27) which provides a list with the impact that the variants between the reference and target genome have on the annotated genes (Suppl Table 1). From the 2,469 total changes, 242 affected protein coding genes (Table 2). From these, 191 genes were affected by substitutions, 129 of them were non-synonymous with two of them introducing a new stop codon. Fifty-one genes were affected by indels from which 18 were in frame and 30 introduced a frameshift, two led to a loss of start codon and one to a 3’ truncation (Table 2).

**Table 2:**
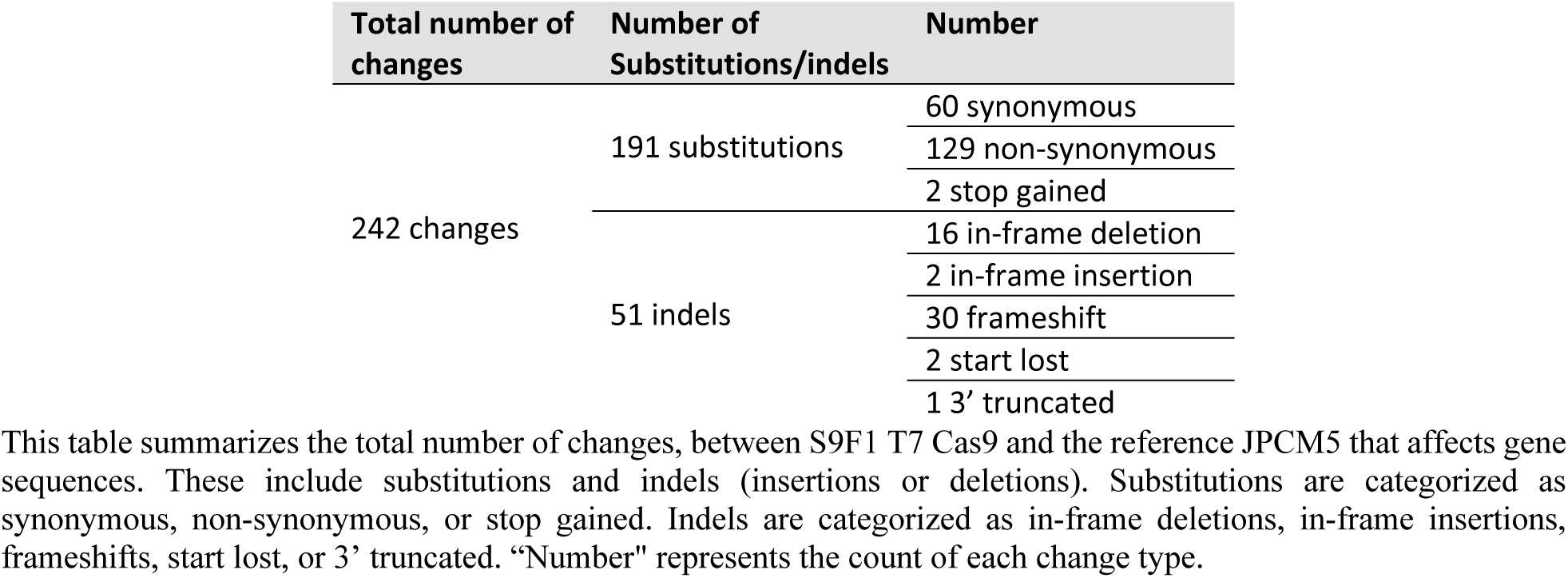
Summary of genome changes of S9F1-Cas9 with effect on gene sequences.

After re-sequencing and annotating the S9F1 T7 Cas9 genome, we integrated the data into LeishGEdit, providing the scientific community with easy access to primer design for CRISPR-Cas9 experiments.

### CRISPR-Cas9 system is efficient in *L. infantum* S9F1

To validate if the T7 Cas9 LeishGEdit system is functional, PF16 was tagged in its endogenous locus, with a 3xHA epitope at the C-terminal end. Immunofluorescence assays revealed as expected a localization of PF16 in the flagellum (Fig 4A). We then generated PF16 knockout parasites by creating gene deletion mutants, where both alleles of PF16 were replaced with antibiotic resistance genes (puromycin and geneticin) in the S9F1 T7 Cas9 background. Null mutants were generated, and gene deletion was confirmed by PCR (Fig. 4B). We assessed the cell growth of both S9F1 T7 Cas9 and S9F1 T7 Cas9-KO PF16, finding no significant difference in growth between the mutant and wild-type strains (Fig. 4C). A motility test revealed that, as expected, the KO-PF16 parasites were non-motile, consistent with the known effects of PF16 deletion in *Leishmania* (11,19,37) (Fig. 4D). These results demonstrate that S9F1 T7 Cas9 is an effective strain for genetic engineering.

**Figure 4:**
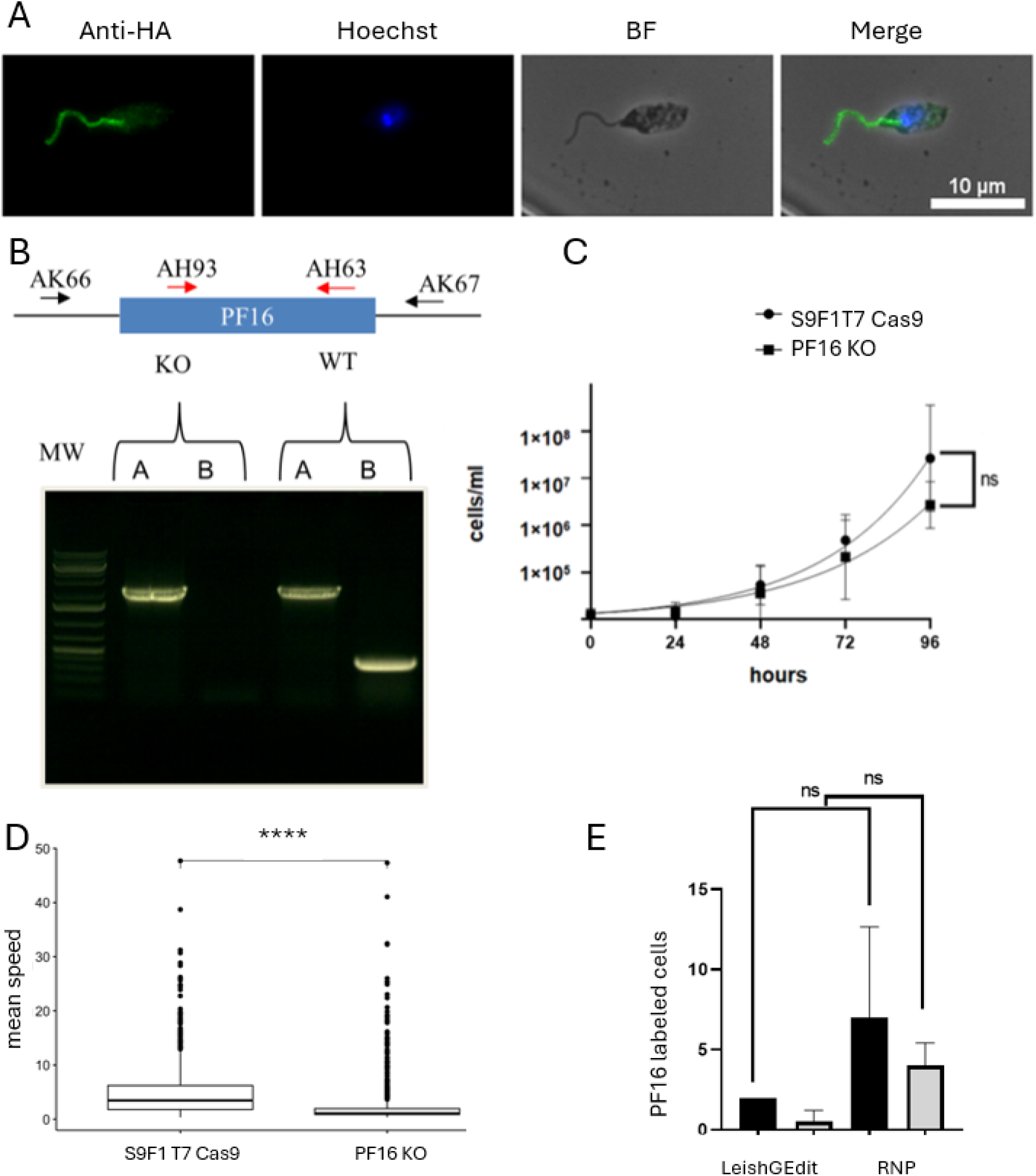
The S9F1 T7 Cas9 LeishGEdit system is functional (A) Flagellum was labeled using C terminus tagging with 3HA of PF16 gene. (B) Verification of the deletion of PF16 by PCR: A: primers AK66/AK67: non-edited 2207bp, edited:2288bp, B: primers AH93/AH63: non-edited 407bp, edited: absence of band. MW: 1Kb Plus gene ruler. (C) Growth curves of S9F1 T7 Cas9 and PF16 KO strains. Statistics: unpaired t-test (two-sided), ns: not significant. (D) Mean speed was measured using an open-source Fiji plugin (Track Mate). Statistical test: student test, ****= p value < 0.0001. (E) Comparison of LeishGEdit and RNP-CRISPR-Cas9 CRISPR-Cas9 strategies. In both cell lines, PF16 was 3’ tagged, marker-free, at its endogenous locus either with an mNG or mCherry cassettes. Around 200 cells were analyzed for each condition, mNG (black histograms), mCherry (grey histograms).

We also wished to validate an alternative strategy in which Cas9 and sgRNAs are synthesized *in vitro* which has already been implemented in trypanosomatids (12,13). The ribonucleoprotein complex is generated *in vitro* and then transfected into parasites. The transient nature of Cas9 expression likely minimizes the risks of toxicity, genome instability, off-target effects, and reduced growth observed in many other cells (35). However, in our experiments, these risks appear to be minimal or nonexistent despite the constitutive expression of Cas9. We performed marker-free tagging of PF16 at its endogenous locus with either mNeonGreen (mNG) or mCherry in parallel with both CRISPR-Cas9 strategies. The experiment was done in triplicates. On average, seven out of 200 cells exhibited flagellar labeling with mNG using the RNP-CRISPR-Cas9 strategy, compared to only two cells labeled with the CRISPR-Cas9 system PCR-based method. Similarly, four out of 200 cells showed flagellar labeling with mCherry using the RNP-CRISPR strategy-Cas9, whereas one cell is labeled with the CRISPR-Cas9 system PCR-based method (Fig. 4E). A trend was therefore identified: with the RNP approach, a higher number of cells were successfully labeled with both mNG and mCherry than with the PCR-based method in the three replicates for each tag. However, these differences did not reach here statistical significance. The RNP approach is easy to perform, as there is no molecular cloning involved, which saves time. However, it is more expensive than the LeishGEdit approach. In the laboratory, we use the RNP approach when we don’t want to use selection markers, or when we want to carry out multiple modifications within a single cell line, such as the editing of large multi-gene families and the introduction of point mutations. However, this approach requires cell cloning and screening of edited cell lines, making it less suitable for rapid, high-throughput screening. In contrast, the LeishGEdit strategy enables complete editing of non-essential genes within a non-clonal population, making it a more efficient choice for large-scale knock-out and tagging screens (26).

To conclude, we determined that S9F1, whose genetic background is that of ITMAP263, was a well-adapted strain for carrying out genetic engineering experiments. This result is supported by the fact that strain ITMAP263 has already been used in a genome-wide CRISPR-Cas9 screen to identify genes involved in drug resistance in *Leishmania infantum* (38). We have successfully re-sequenced and annotated the S9F1 T7 Cas9 genome, integrating this data into LeishGEdit. This provides the scientific community with accessible genomic information and gene annotations to aid in designing CRISPR-Cas9 experiments. Our next goal is to make this strain widely available to support pathogenesis studies of visceral leishmaniasis.

## AUTHOR CONTRIBUTIONS

RT: Investigation, Methodology, Software, Formal Analysis, Visualization, Writing – Original Draft Preparation, MSC: Investigation, TB: Software, Writing –Review & Editing, NK: Resources (strains), Investigation, GP: Investigation, Software and Formal Analysis (live microscopy, Trackmate), DJ: Resources (NGS sequencing), Writing – Review & Editing, BV: Resources (strains), Writing – Review & Editing, BB: Methodology, Software, Data Curation, Writing – Original Draft Preparation-– Review & Editing and YS: Conceptualization, Project Administration, Funding Acquisition, Methodology, Data Curation, Supervision, Validation, Writing – Original Draft Preparation-– Review & Editing

## DATA AVAILABILITY

The whole genome sequencing data (FASTQ files) from this study have been submitted to the European Nucleotide Archive under the accession number PRJEB85295.

## ACKNOWLEDGEMENTS

YS received a grant and RT and BB were funded by the French LabEx ParaFrap (ANR-11-LABX-0024). We would like to thank the EuPathDB database (https://veupathdb.org/veupathdb/app) which was essential for the completion of this study. We would like to thank the National Reference Center for Leishmaniasis and the BRC-*Leishmania* for providing us with *L. infantum* LEM6696 (MCAN/ES/98/LLM-877) and *L. donovani* LEM703 (MHOM/IN/80/DD8), Gerald Späth (Unité Ld1S-Bob, Eva Gluenz (Molecular Cell biology of *Leishmania,* Institute of Cell Biology, Universität Bern) for providing us with *L. mexicana* (MNYC/BZ/62/M379) and with *L. mexicana* T7 Cas9 derived from the same genetic background by transfection of the pTB007 plasmid. The LACK antibody was a kind gift from Eric Prina (Unité Parasitologie moléculaire et Signalisation, Institut Pasteur). Finally, we would like to extend our warmest thanks to Eva Gluenz (Molecular Cell biology of *Leishmania,* Institute of Cell Biology, Universität Bern) for agreeing to integrate our data into LeishGedit, an essential prerequisite for the distribution of our strain to the community.

## REFERENCES

1. Global leishmaniasis update, 2006–2015: a turning point in leishmaniasis surveillance. Wkly Epidemiol Rec. 2017 Sep 22;92(38):557–65.

2. Sunter J, Gull K. Shape, form, function and Leishmania pathogenicity: from textbook descriptions to biological understanding. Open Biol. 2017 Sep;7(9):170165.

3. Akhoundi M, Kuhls K, Cannet A, Votýpka J, Marty P, Delaunay P, et al. A Historical Overview of the Classification, Evolution, and Dispersion of Leishmania Parasites and Sandflies. PLoS Negl Trop Dis. 2016 Mar 3;10(3):e0004349.

4. Torres-Guerrero E, Quintanilla-Cedillo MR, Ruiz-Esmenjaud J, Arenas R. Leishmaniasis: a review. F1000Res. 2017 May 26;6:750.

5. Costa CHN, Chang KP, Costa DL, Cunha FVM. From Infection to Death: An Overview of the Pathogenesis of Visceral Leishmaniasis. Pathogens. 2023 Jul 24;12(7):969.

6. Reis-Cunha JL, Grace CA, Ahmed S, Harnqvist SE, Lynch CM, Boité MC, et al. The global dispersal of visceral leishmaniasis occurred within human history [Internet]. bioRxiv; 2024 [cited 2025 Feb 13]. p. 2024.10.30.621037. Available from: https://www.biorxiv.org/content/10.1101/2024.10.30.621037v1

7. Dereure J, El-Safi SH, Bucheton B, Boni M, Kheir MM, Davoust B, et al. Visceral leishmaniasis in eastern Sudan: parasite identification in humans and dogs; host-parasite relationships. Microbes Infect. 2003 Oct;5(12):1103–8.

8. Pratlong F, Dereure J, Bucheton B, El-Saf S, Dessein A, Lanotte G, et al. Sudan: the possible original focus of visceral leishmaniasis. Parasitology. 2001 Jun;122(Pt 6):599–605.

9. Sollelis L, Ghorbal M, Macpherson CR, Martins RM, Kuk N, Crobu L, et al. First efficient CRISPR-Cas9-mediated genome editing in Leishmania parasites. Cellular Microbiology. 2015;17(10):1405–12.

10. Yagoubat A, Corrales RM, Bastien P, Lévêque MF, Sterkers Y. Gene Editing in Trypanosomatids: Tips and Tricks in the CRISPR-Cas9 Era. Trends in Parasitology. 2020 Sep 1;36(9):745–60.

11. Beneke T, Madden R, Makin L, Valli J, Sunter J, Gluenz E. A CRISPR Cas9 high-throughput genome editing toolkit for kinetoplastids. R Soc Open Sci. 2017 May;4(5):170095.

12. Soares Medeiros LC, South L, Peng D, Bustamante JM, Wang W, Bunkofske M, et al. Rapid, Selection-Free, High-Efficiency Genome Editing in Protozoan Parasites Using CRISPR-Cas9 Ribonucleoproteins. mBio. 2017 Nov 7;8(6):e01788–17.

13. Asencio C, Hervé P, Morand P, Oliveres Q, Morel CA, Prouzet-Mauleon V, et al. Streptococcus pyogenes Cas9 ribonucleoprotein delivery for efficient, rapid and marker-free gene editing in Trypanosoma and Leishmania. Mol Microbiol. 2024 Jun;121(6):1079–94.

14. Berens RL, Brun R, Krassner SM. A simple monophasic medium for axenic culture of hemoflagellates. J Parasitol. 1976 Jun;62(3):360–5.

15. Bates PA. Complete developmental cycle of Leishmania mexicana in axenic culture. Parasitology. 1994 Jan;108 ( Pt 1):1–9.

16. Campos M, Gomes C, Souza A, Lainson R, Corbett C, Silveira F. In vitro infectivity of species of Leishmania (Viannia) responsible for American cutaneous leishmaniasis. Parasitology research. 2008 Jun 1;103:771–6.

17. Corrales RM, Vaselek S, Neish R, Berry L, Brunet CD, Crobu L, et al. The kinesin of the flagellum attachment zone in Leishmania is required for cell morphogenesis, cell division and virulence in the mammalian host. PLoS Pathog. 2021 Jun 18;17(6):e1009666.

18. Tinevez JY, Perry N, Schindelin J, Hoopes GM, Reynolds GD, Laplantine E, et al. TrackMate: An open and extensible platform for single-particle tracking. Methods. 2017 Feb 15;115:80–90.

19. Martel D, Beneke T, Gluenz E, Späth GF, Rachidi N. Characterisation of Casein Kinase 1.1 in Leishmania donovani Using the CRISPR Cas9 Toolkit. Biomed Res Int. 2017;2017:4635605.

20. Mougneau E, Altare F, Wakil AE, Zheng S, Coppola T, Wang ZE, et al. Expression cloning of a protective Leishmania antigen. Science. 1995 Apr 28;268(5210):563–6.

21. Dean S, Sunter JD, Wheeler RJ. TrypTag.org: A Trypanosome Genome-wide Protein Localisation Resource. Trends in Parasitology. 2017 Feb;33(2):80–2.

22. Halliday C, Billington K, Wang Z, Madden R, Dean S, Sunter JD, et al. Cellular landmarks of Trypanosoma brucei and Leishmania mexicana. Molecular and Biochemical Parasitology. 2019 Jun 1;230:24–36.

23. Martin M. Cutadapt removes adapter sequences from high-throughput sequencing reads. EMBnet.journal. 2011 May 2;17(1):10–2.

24. Bolger AM, Lohse M, Usadel B. Trimmomatic: a flexible trimmer for Illumina sequence data. Bioinformatics. 2014 Aug 1;30(15):2114–20.

25. Danecek P, Bonfield JK, Liddle J, Marshall J, Ohan V, Pollard MO, et al. Twelve years of SAMtools and BCFtools. Gigascience. 2021 Feb 16;10(2):giab008.

26. Li H, Handsaker B, Wysoker A, Fennell T, Ruan J, Homer N, et al. The Sequence Alignment/Map format and SAMtools. Bioinformatics. 2009 Aug 15;25(16):2078–9.

27. Shumate A, Salzberg S. LiftoffTools: a toolkit for comparing gene annotations mapped between genome assemblies. F1000Res. 2024 Apr 29;11:1230.

28. Beneke T, Gluenz E. Bar-seq strategies for the LeishGEdit toolbox. Molecular and Biochemical Parasitology. 2020 Sep 1;239:111295.

29. Aellig S, Billington K, Damasceno JD, Davidson L, Dobramysl U, Etzensperger R, et al. LeishGEM: genome-wide deletion mutant fitness and protein localisations in *Leishmania*. Trends in Parasitology. 2024 Aug 1;40(8):675–8.

30. Denise H, Poot J, Jiménez M, Ambit A, Herrmann DC, Vermeulen AN, et al. Studies on the CPA cysteine peptidase in the Leishmania infantum genome strain JPCM5. BMC Molecular Biology. 2006 Nov 13;7(1):42.

31. Bianchi L, Rondanelli EG, Carosi G, Gerna G. Endonuclear Mitotic Spindle in the Leptomonad of Leishmania tropica. The Journal of Parasitology. 1969 Oct;55(5):1091.

32. Triemer RE, Fritz LM, Herman R. Ultrastructural features of mitosis inLeishmania adleri. Protoplasma. 1986 Jun 1;134(2):154–62.

33. Liu B, Liu Y, Motyka SA, Agbo EEC, Englund PT. Fellowship of the rings: the replication of kinetoplast DNA. Trends in Parasitology. 2005 Aug 1;21(8):363–9.

34. Ploubidou A, Robinson DR, Docherty RC, Ogbadoyi EO, Gull K. Evidence for novel cell cycle checkpoints in trypanosomes: kinetoplast segregation and cytokinesis in the absence of mitosis. Journal of Cell Science. 1999 Dec 15;112(24):4641–50.

35. Ambit A, Woods KL, Cull B, Coombs GH, Mottram JC. Morphological Events during the Cell Cycle of Leishmania major ▿. Eukaryot Cell. 2011 Nov;10(11):1429–38.

36. Wheeler RJ, Gluenz E, Gull K. The cell cycle of Leishmania: morphogenetic events and their implications for parasite biology. Mol Microbiol. 2011 Feb;79(3):647–62.

37. Yagoubat A, Crobu L, Berry L, Kuk N, Lefebvre M, Sarrazin A, et al. Universal highly efficient conditional knockout system in Leishmania, with a focus on untranscribed region preservation. Cell Microbiol. 2020;22(5):e13159.

38. Queffeulou M, Leprohon P, Fernandez-Prada C, Ouellette M, Mejía-Jaramillo AM. CRISPR-Cas9 high-throughput screening to study drug resistance in Leishmania infantum. mBio. 15(7):e00477–24.

